# Machine learning based classification of deep brain stimulation outcomes in a rat model of binge eating using ventral striatal oscillations

**DOI:** 10.1101/241794

**Authors:** Wilder T. Doucette, Lucas Dwiel, Jared E. Boyce, Amanda A. Simon, Jibran Y. Khokhar, Alan I. Green

**Author notes:** **Corresponding Author:** Wilder T. Doucette, MD, PhD 1 Medical Center Drive, Lebanon, NH 03756.

## Abstract

Neuromodulation-based interventions continue to be evaluated across an array of appetitive disorders but broader implementation of these approaches remains limited due to variable treatment outcomes. We hypothesize that individual variation in treatment outcomes may be linked to differences in the networks underlying these disorders. Here, Sprague-Dawley rats received deep brain stimulation separately within each nucleus accumbens (NAc) sub-region (core and shell) using a within-animal crossover design in a rat model of binge eating. Significant reductions in binge size were observed with stimulation of either target but with significant variation in effectiveness across individuals. When features of local field potentials (LFPs) recorded from the NAc were used as predictors of the pre-defined stimulation outcomes (response or non-response) from each rat using a machine-learning approach (lasso), stimulation outcomes could be predicted with greater accuracy than expected by chance (effect sizes: core = 1.13, shell = 1.05). Further, these LFP features could be used to identify the best stimulation target for each animal (core vs. shell) with an effect size = 0.96. These data suggest that individual differences in underlying network activity may contribute to the variable outcomes of circuit based interventions and that measures of network activity have the potential to individually guide the selection of an optimal stimulation target and improve overall treatment response rates.

## Introduction

Brain stimulation has demonstrated the potential to improve symptoms in Parkinson’s disease, depression and obsessive-compulsive disorder, yet highly variable treatment outcomes (especially common in psychiatric disorders) indicate that the full potential of brain stimulation is not being met [1–3]. The majority of these studies evaluate the treatment outcomes of a single brain target despite pre-existing evidence supporting the potential of other stimulation targets [2, 4–6]. With these constraints, treatment outcome improvements have mostly been achieved to date through more stringent inclusion criteria and improved precision in modulating the intended brain target [7–9]. Another potential avenue to improve treatment outcomes for a specific disorder could be achieved through the personalization of target selection. This approach was pioneered by cancer biologists that used tumor immunoprofiling to personalize chemotherapy and it remains unknown if personalization of target selection for neuromodulation-based treatments has a similar potential to improve treatment outcomes in neuropsychiatric diseases including disorders of appetitive behavior.

Clinical studies that used invasive or non-invasive stimulation in disorders of appetitive behavior (e.g., addiction, binge eating and obesity), have demonstrated the potential of targeting an array of different brain areas but also demonstrated considerable treatment response heterogeneity across individuals [5, 10–14]. The pre-clinical literature on deep brain stimulation (DBS), while also encouraging for appetitive disorders, reveals considerable outcome variation resulting from the targeting of different brain regions across studies. In addition, most studies report only population-based effects, masking the problem of variation across individuals [15–17].

In this study, we used an established rat model of binge eating to produce binge-like feeding behavior [18–20], Similar rodent models of binge eating have resulted in weight gain[18], compulsive feeding behavior[21, 22] and increased impulsivity[23] thus displaying problems in overlapping psychological domains to patients with binge eating disorder. It is important to acknowledge however, that this is merely a pre-clinical approximation of the clinical condition and many successful pharmacologic trials using this rodent/rat model have failed to translate clinically with the exception of lisdexamfetamine [24, 25]. Using this pre-clinical model of binge eating we have previously shown variation in individual rat outcomes receiving deep brain stimulation targeting the nucleus accumbens core with about 60% of rats displaying a significant reduction in binge size with stimulation [26]. When non-invasive, repetitive transcranial magnetic stimulation was targeted to a related area of the reward circuit in patients with binge eating, the frequency of binges decreased in 18 of 28 subjects (~60%) [27]. While the primary outcome in clinical and pre-clinical studies is necessarily different (frequency of binges vs. size of binges) this rat model of binge eating could provide insight into the source of stimulation outcome variability and provide a model to explore the potential feasibility and benefit of personalized target selection for stimulation-based interventions.

We theorize that individual variation in brain stimulation outcomes targeting a specific brain region may be linked to individual differences in the networks underpinning the symptom of interest (e.g., binge eating) [27]. It follows that measures of relevant network activity would be able to predict brain stimulation outcomes at a given brain target or be used to individualize the choice between potentially viable targets. This study was designed to compare the treatment efficacy of stimulation targeted to either the nucleus accumbens (NAc) core or shell, two regions with known differences in anatomic and functional connectivity and differential functional roles across an array of reward related behaviors [28, 29]. This study replicated our previous treatment outcome variance with NAc core stimulation [26] and extended the results to assess whether similar variation in treatment outcomes occurs with NAc shell stimulation (previously reported by Halpern et al. to be effective in a mouse model of binge eating) [30]. It was then determined whether a correlation existed between individual stimulation outcomes and either corresponding performance on reward-related behaviors, local field potential recordings from the ventral striatum, or electrode localization within each NAc sub-region.

## Methods and Materials

### Animals and Surgery

Male Sprague-Dawley rats were purchased from Charles River *(Shrewsbury, MA)* at 60 days of age and individually housed on a reverse 12 hour light/dark schedule with house chow and water available *ad libitum*. Following habituation to the animal facility, rats were implanted with a custom electrode array that targeted both the NAc core and shell bilaterally, according to the following coordinates relative to bregma: 1.6 mm anterior; ± 1 and 2.5 mm lateral; and 7.6 mm ventral. Animals were excluded from analysis if later histologic examination revealed electrode location outside of the NAc core or shell. All experiments were carried out in accordance with the NIH Guide for the Care and Use of Laboratory Animals (NIH Publications No. 80–23) revised in 1996 and approved by the Institutional Animal Care and Use Committee at Dartmouth College.

### Binge Eating Paradigm

Following recovery from surgery (~1 week), rats began a paradigm of limited access to a palatable high-fat, high-sugar diet (“sweet-fat diet”), which contained 19% protein, 36.2% carbohydrates, and 44.8% fat by calories and 4.6 kcal/g (Teklad Diets 06415, *South Easton, MA)* as previously described [18]. The sweet-fat diet was provided to the rats in addition to house chow and water within stimulation chambers for 2 hour sessions, with 4–5 sessions per week (irregular schedule). Following 16–20 sessions the rats were consuming a stable and significant amount of sweet-fat food during each session (mean = 54% of their daily caloric intake ± 12% [1 standard deviation]). This “binge-like” feeding has been shown to result in more significant weight gain than is observed with continuous access to the same diet -- as is used in models of diet-induced obesity [18]. Prior work has also demonstrated that chronic, irregular, limited access to palatable food can result in compulsive feeding behavior[21, 22] and increased impulsivity [23]. Palatable sweet-fat and regular house chow consumption was measured during all limited access sessions.

### Stimulation

To deliver stimulation, a current-controlled stimulator *(PlexStim, Plexon, Plano, TX)* was used to generate a continuous train of biphasic pulses. The output of the stimulator (current and voltage) was verified visually for each rat before and after each stimulation session using a factory-calibrated oscilloscope *(TPS2002C, Tektronix, Beaverton, OR)*. Stimulation was initiated immediately before animals had access to the sweet-fat food and turned off at the completion of the 2 hour session.

### Overall Design

Experiment 1 (N=8 rats) was used to determine the optimal stimulation parameters to reduce binge size using our custom electrode arrays targeting the NAc core or shell. Experiment 2 used a crossover design in a separate cohort of 9 rats to test DBS targeting the NAc core or shell with the optimized stimulation parameters from Experiment 1. Lastly, rats from Experiment 1 and 2 that had received the optimized stimulation parameters in both NAc targets and remained in good health (N=12) continued on to Experiment 3 and underwent behavioral and electrophysiological characterization (Figure 1A).

**Figure 1.**
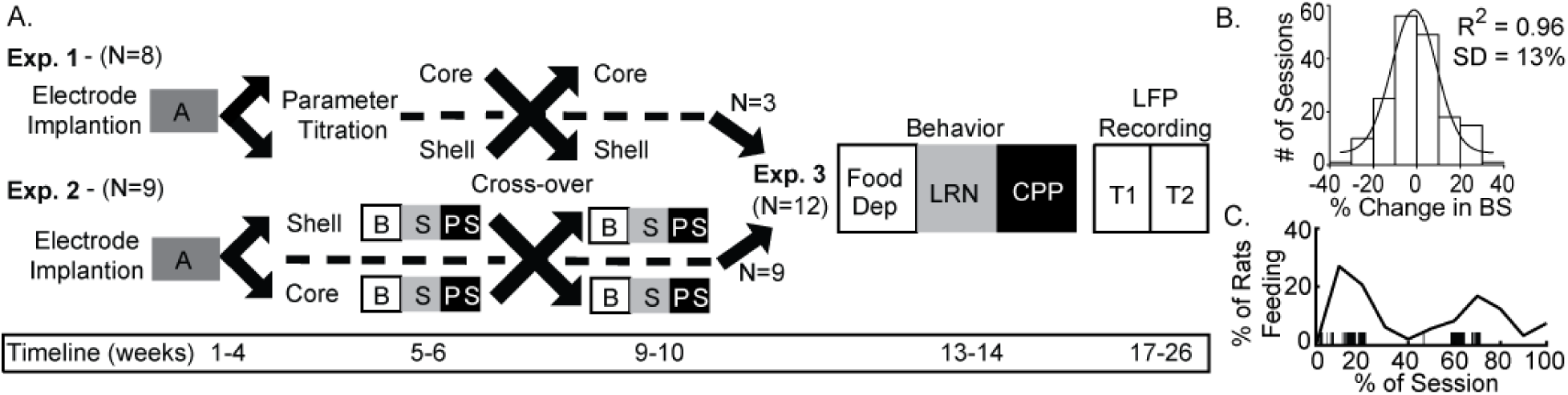
Experimental design and timeline with population data used to define significant change in binge size from baseline. **A**. Experimental design for Experiments 1–3 with timeline shown at bottom. A - acquisition of stable binge size following chronic irregular limited access and randomization to initial stimulation target, B - baseline sessions, S - stimulation sessions, PS - post-stimulation sessions, Food Dep - food deprived binge session, LRN - locomotor response to novelty, CPP - conditioned place preference, LFP Recording - local field potential recording at two timepoints (T1 and T2). **B**. Population baseline data (3 sessions per animal, N=36 animals) was used to determine an *a priori* definition of a significant change from baseline binge size (BS). Distribution of binge size variance across baseline sessions was fit to a normal distribution with R^2^ fit shown (1 standard deviation [SD] = 13% change from baseline average). **C**. The percentage of animals engaged in feeding behavior through a normalized binge session had a bimodal distribution. Vertical black lines under the curve provide an individual example of all of the feeding epochs from a single animal through a binge session.

## Experiment 1 - Identifying optimal stimulation parameters

To identify the optimal stimulation parameters for our custom arrays within the target brain structures (NAc core and shell) for altering feeding behavior we tested an array of published stimulation intensities (range: 150 to 500 µA) and electrode contact configurations (monopolar vs. bipolar). These permutations alter the size and shape of the electric field and the resulting effect that stimulation has on binge eating. Thus, custom electrodes were implanted in the NAc core and shell bilaterally in a cohort of rats (N=8). Using a simple randomization, rats were divided into two groups for a crossover design with different initial stimulation targets (core or shell). Animals were then trained in the binge eating paradigm until a stable baseline of sweet-fat food intake was established (15–20 sessions over 3–4 weeks) before stimulation sessions were initiated. Stimulation current was increased for each subsequent session, starting at 150 µA and progressing to 500 μA in a bipolar configuration (between two wires within the target, separated by ~1mm in the dorsal-ventral plane), and then from 150 μA to 300 μA in a monopolar configuration (between one wire in the target and a skull screw over lambda). Animals then re-established their pre-stimulation baseline of sweet-fat diet consumption in the absence of stimulation. Following a return to baseline rats initiated stimulation sessions in the other NAc target with subsequent titration of stimulation parameters across multiple sessions (Figure 1A).

## Experiment 2 - Testing NAc core vs. shell stimulation using fixed stimulation parameters

Experiment 1 identified stimulation parameters that were similarly effective in either the NAc core or shell-bipolar stimulation at 300 μA or monopolar stimulation at 200 μA. We elected to use monopolar stimulation (biphasic, 90 µsec pulse width, 130 Hz, 200 μA) as it produced a lower charge density at the electrode surface which decreases the probability of neuronal injury [31]. In a new cohort of rats (N=9) electrodes were implanted and rats were randomized to receive initial stimulation in either the NAc core or shell using a simple randomization. After a stable baseline of sweet-fat diet consumption was established during limited access sessions (following 15–20 sessions), rats received 3 sessions of stimulation followed by 3 sham post-stimulation sessions. Animals then entered a **2 week washout phase** to re-establish baseline prior to crossover and stimulation in the other target (Figure 1A).

## Data Analysis

### Experiment 1 data analysis

In order to evaluate experiment 1 data an *a priori* definition of what a meaningful DBS induced change in binge size was required. This was accomplished by pooling baseline binge eating data from multiple cohorts to characterize variation in baseline binge size within the population (36 rats, 3 baseline sessions per rat, 108 total baseline observations). The data came from all of the animals in this study, a previously published study [26], and unpublished data. Each observation was recorded as the percent change from that rats average baseline binge size. This “normalized variance” was done to account for the known variation between animals in their average binge size at baseline. This session to session normalized variation in binge size was found to be normally distributed, centered at 0% change with a standard deviation of 13% (Figure 1B). Thus, for Experiment 1, if an animal’s binge size during a stimulation session was greater or less than 26% (2 standard deviations) of its average baseline binge size it was considered a meaningful change induced by stimulation.

### Experiment 2 data analysis

#### Population-based analysis

We used standard population-based analysis for repeated observations (repeated measures analysis of variance, RMANOVA) and included 3 sessions of baseline, stimulation and post-stimulation data from each animal. Each stimulation target was analyzed independently as there were no significant differences in binge size between the baseline periods on either side of the crossover. Session number (1–3) and session type (baseline, stimulation, and post-stimulation) were used as independent variables. When the analysis indicated that differences existed between session types, post-hoc pair-wise comparisons between groups were made and multiple comparisons were corrected using Bonferroni with a corrected significance level set at p≤0.05.

#### Individual-based analysis

Given our previously observed individual variation in DBS response we characterized each animal as either a responder or non-responder to stimulation in each target using an *a priori* definition. These individual stimulation response categories were then used for subsequent correlation with reward-related behavior and electrophysiologic recordings. Individual rats were classified as either non-responders [NR] or responders [R] to stimulation at each target based on the criteria used in Experiment 1 (greater than a 2 SD or 26% change in binge size from each animal’s baseline average) and this change had to be observed in all three stimulation sessions for a given target.

## Experiment 3 - Behavioral and electrical characterization (without stimulation)

All rats from Experiment 2 (N=9) and those rats from Experiment 1 tested with the stimulation parameters chosen for Experiment 2 in both targets (N=3) were included in Experiment 3 (N=12). These animals underwent subsequent behavioral and electrical characterization starting two weeks after the conclusion of Experiment 1 or 2. All rats underwent behavioral testing followed by another 2 week washout and then electrophysiology, **all without stimulation** (Figure 1A).

### *Reward-related behavior* (order of testing)

To determine if variation in reward-related behavior could capture the underlying network differences that may be responsible for the variation in DBS outcomes, 3 reward-related behaviors were assessed. Behavioral outcomes were compared between NR and R groups for each DBS target using a two-way t-test. A significance threshold of p<0.05 was used to screen for behaviors with potential correlation to stimulation outcomes.

#### Increased sweet-fat diet intake with food deprivation (1)

Food deprivation (24 hours) was used to push the energy homeostasis system towards an orexigenic state. Individual variation in the resultant changes in binge size from baseline was measured. Thus, the primary outcome was the percent change in binge size from each rat’s baseline average to that observed following food deprivation.

#### Locomotor response to novelty (2)

Locomotor response to novelty was chosen because of previous correlations between variation in this behavior (high and low responders) and a sensation-seeking behavioral phenotype linked to a higher risk for developing disorders of appetitive behavior [32, 33]. Briefly, rats were placed in a 1.5 ft × 3 ft black plastic chamber that was novel to the animal and allowed to freely explore for 50 minutes while video was recorded. Video files were analyzed offline using automated contrast-based tracking (Cineplex software, *Plexon, Plano, TX)* to calculate the distance traveled (primary outcome).

#### Conditioned place preference (CPP) (3)

CPP was assessed due to the known involvement of the NAc in CPP [34]. We used an established 2-chamber biased design paradigm, pairing the sweet-fat food with the individual animal’s non-preferred chamber and regular house chow with the preferred chamber (30 minute pairing, 1 pairing per day, alternating between the 2 chambers for 4 days) [35, 36]. Baseline and test sessions (15 minutes) were video recorded and automatically scored using contrast-based tracking to assess time spent in each chamber. The primary outcome was the change in the percentage of time spent in the initially non-preferred chamber (paired with sweet-fat diet).

### Local field potential (LFP) recording

We recorded local field potential (LFP) activity bilaterally from the NAc core and shell of each animal to assess if variation of intrinsic network characteristics in the absence of stimulation correlated with stimulation outcomes. Rats were tethered in a neutral chamber through a commutator to a Plexon data acquisition system while time-synchronized video was recorded *(Plexon, Plano, Tx)* for offline analysis. Using the video, rest intervals were manually identified as extended periods of inactivity and only recordings from these intervals were used in the analysis. We used well-established frequency ranges from the rodent literature and standard LFP signal processing to characterize the power spectral densities (PSDs) within, and coherence between brain regions (bilateral NAc core and shell) for each animal using custom code written using Matlab R2015b [37–39] (Supplemental Methods). Each rat recording session produced 60 LFP features: 24 measures of power (6 frequency bands X 4 brain locations) and 36 measures of coherence (6 frequency bands X 6 possible location pairs, Figure 5A and B).

### Linking ventral striatal activity to stimulation outcomes

As there were many more predictor variables than number of animals we employed a machine learning approach to determine if there was information within the LFP signals that correlated with stimulation outcomes. We used a penalized regression method, lasso, to reduce the dimensionality by removing LFP features that contained no information or redundant information and thereby extract a combination of LFP features that most accurately described the observed variation in stimulation outcomes. The Matlab package *Glmnet* was used to implement the lasso using a cross-validation scheme with 100 repetitions for each model (Core R vs. NR, Shell R vs. NR, and Core vs. Shell). For the Core vs. Shell model, each animal’s optimal stimulation target was defined as the stimulation target that produced the largest average reduction in binge size (rats without a significant reduction were excluded). The accuracy of the models is reported as the average cross-validated accuracy. In order to determine if the achieved accuracies were meaningfully better then chance, the entire process stated above was repeated for ten random permutations of the data for each model type. The permutations randomized the relationship between the binary stimulation outcomes (R=1, NR=0) or optimal target assignment (Core =1, Shell=0) with the individual rat LFP feature sets to maintain the overall structure of the data but permute the relationship of dependent to independent variables. The distribution of accuracies from the observed data was then compared to the distribution from the permuted data using the Mann-Whitney U test and then the U test statistic was converted into a Cohen’s d effect size.

If the lasso indicated that information existed in the LFP signal, a subsequent investigation of each LFP feature was carried out to determine which features contained the most information. For this, univariate logistic models were built using each of the LFP features to classify: 1) core responses; 2) shell responses; or 3) core or shell as the best stimulation target for each animal. For the logistic models, an exhaustive leave-one-out cross-validation was used in order to obtain a distribution of accuracies and the mean accuracy from these distributions is reported in Table 1 for the top 5 features from each model type.

**Table 1.**
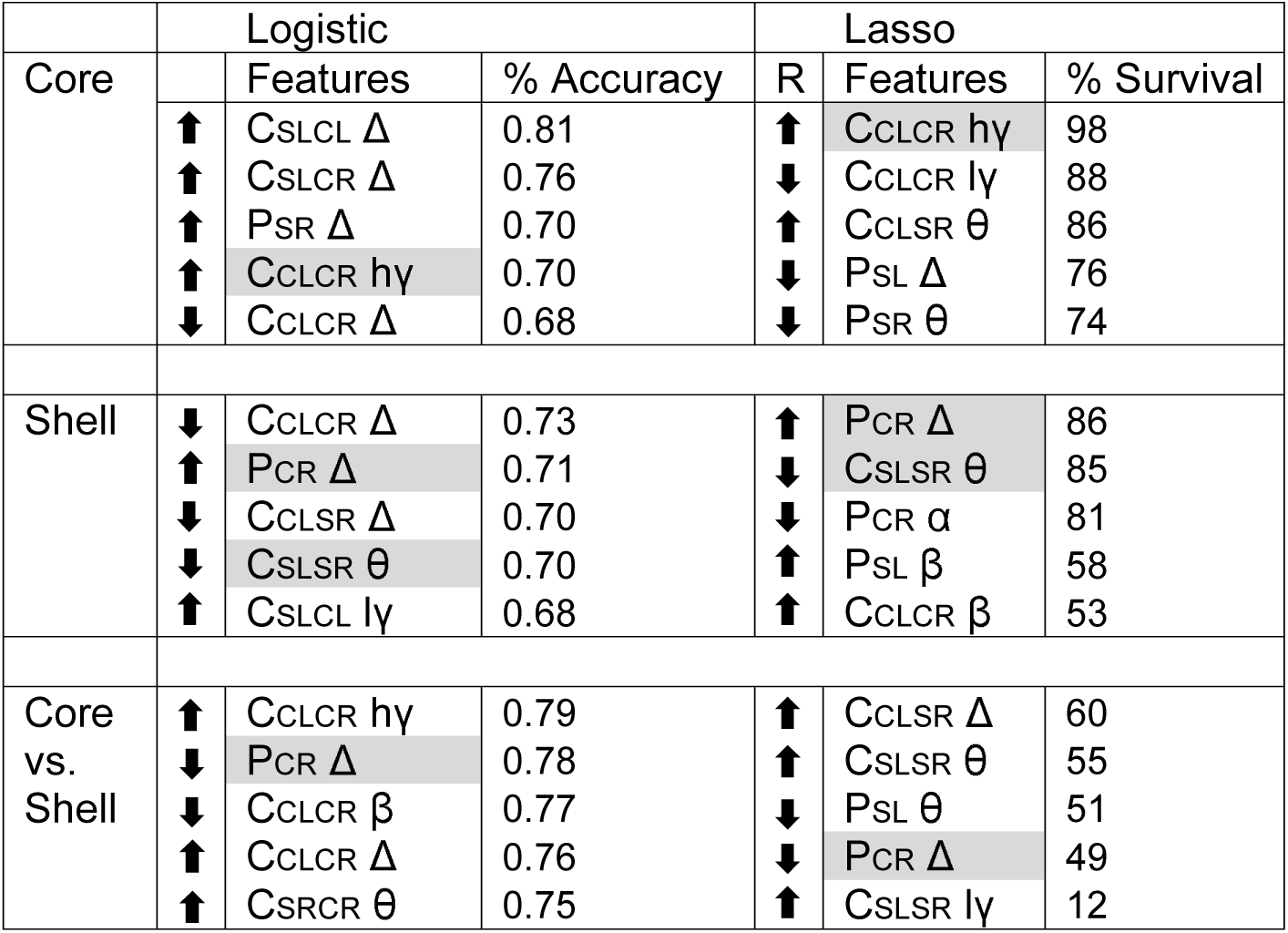
Top 5 LFP Features for Each Model Type. The top 5 local field potential features used in single predictor (logistic) and multi-predictor (lasso) models of NAc core and shell stimulation outcomes. Features are described by location (Core Left -CL, Core Right -CR, Shell Left -SL, and Shell Right -SR) and frequency band (delta -△, theta -θ, alpha -α, beta -β, low gamma -lγ, and high gamma -hγ). Power features are represented with location and frequency band (e.g., P_SR_ △) and coherence features are represented with location pairs and frequency band (e.g., C_SLCL_ △). Logistic features were ranked by the average % accuracy of the single variable logistic model using leave one out cross-validation. Lasso features were ranked by how frequently they were used in the lasso models from 100 iterations of cross-validation (% survival). The top five features that were common across logistic and lasso models for a given classification type (e.g., core response [R] vs. non-response) are highlighted in grey. Arrows to the left of the LFP feature indicate whether higher (up) or lower (down) LFP feature values increased the probability of a DBS response (R), or in the Core vs. Shell model the direction that increased the likelihood that Core is the better target for that animal.

### Verification of electrode placement

At the conclusion of all experiments rats were euthanized and the brains were removed, prepared for cryostat sectioning, and then mounted and stained (thionine) for histologic analysis of electrode placement [26]. All animals included in the results had electrodes located within the target structure (Figure 4C).

## Results

### Experiment 1 - Identifying optimal stimulation parameters

Figure 2A summarizes the outcome of stimulation in the NAc core with significant reductions in food intake observed with a bipolar configuration (300 µA) in 3/8 animals and with monopolar configuration (200–300 µA) in 4/8 animals. Figure 2B summarizes the outcomes of stimulation of the NAc shell with significant reductions in food intake observed in a subset of animals in bipolar and monopolar configurations. Interestingly, a subset of the shell-stimulated animals had significant increases in food intake at higher stimulation intensities. An example of an individual rat’s food intake across tested stimulation parameters in the NAc core and shell is shown in Figure 2C. This rat had significant reductions in food intake during stimulation in the NAc shell at bipolar 300 μA and monopolar 200 μA with no significant food intake changes with core stimulation (shell only). Figure 2D illustrates the entire cohort’s individual response profiles.

**Figure 2.**
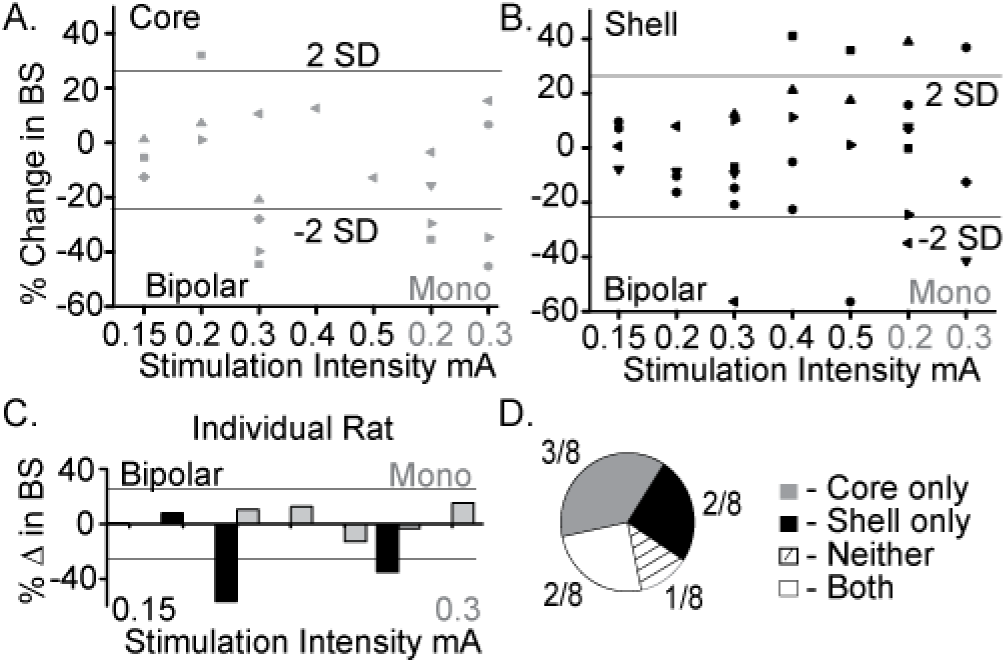
Optimal stimulation parameters were identified that could reduce binge size (BS) using the electrode arrays targeting the NAc core and shell. **A**. Titration of stimulation parameters in NAc core reveals bipolar 300µA and monopolar 200µA are both effective and roughly equivalent. Bipolar (black) and monopolar (Mono, grey) stimulation configurations with corresponding current intensities shown on x-axis. **B**. Titration of stimulation parameters in NAc shell showing similar effective parameters. **C**. Example of a single rat’s stimulation response profile illustrating a shell only responder (core - grey; shell - black). Horizontal lines illustrate ± 2 standard deviations (± 26%). **D**. Distribution of stimulation response profiles for this cohort showing that 5/8 animals resonded to only one of the two stimulation targets.

As demonstrated by the example rat, many animals responded to stimulation in only one of the two NAc sub-regions, despite testing across a range of stimulation parameters. Overall, this cohort of animals helped identify a stimulation configuration *([monopolar]* and parameters *[130 Hz, 90 µsec pulse width, and 200 μA])* for the custom arrays that was capable of decreasing food intake when targeting either the NAc core or shell.

### Experiment 2 - Testing NAc core vs. shell stimulation using optimized stimulation parameters

Figure 3A shows the population outcomes for this cohort (N=9). Using standard population statistics (RMANOVA), a main effect for session type (baseline, stimulation, post-stimulation) was observed in the shell stimulation set (F(1,8) = 8.171, P = 0.02) and in the core stimulation set (F(1,7) = 3.772, P = 0.05). In order to determine which sessions were different, post-hoc pairwise comparisons with Bonferroni adjustment showed a significant difference between the baseline sessions and each stimulation session (p<0.05) but not between the baseline sessions and the post-stimulation sessions.

**Figure 3.**
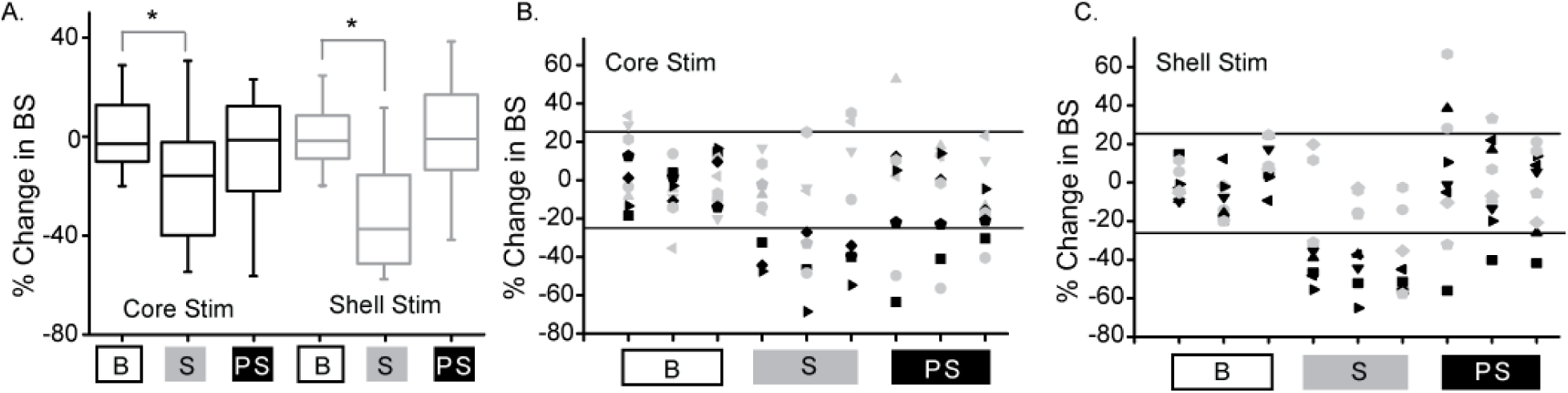
Deep brain stimulation targeted to either the NAc core or shell produces significant reductions in binge size using population-based analysis but with clear individual responders and non-responders. **A**. Population-based analysis (RMANOVA) with post-hoc evaluation revealed a significant difference between baseline (B) and stimulation (S) sessions but not between baseline and post-stimulation (PS) sessions with either core (black) or shell (grey) targeted stimulation (* p≤0.05, boxplots - 95% CI). **B**. Individual rat responses to core stimulation with responders (black, 4/9) and non-responders (grey, 5/9). Horizontal lines illustrate ± 2 standard deviations (± 26%). **C**. Individual rat responses to shell stimulation with responders (black, 5/9) and non-responders (grey, 4/9).

To determine which rats responded to NAc core and shell stimulation, an *a priori* definition of a significant stimulation response was used -- greater than a 26% change from baseline average in all 3 stimulation sessions. The individual responses to NAc core and shell stimulation are shown in Figure 3B and C respectively, with significant individual responders shown in black and non-responders shown in grey. In this cohort, 5/9 rats responded to shell stimulation, 4/9 rats responded to core stimulation and 5/9 rats responded to stimulation in only one of the two targets. Overall (Experiment 1 and 2) 10/17 rats (~60%) responded to only one of the two stimulation targets highlighting the need for individualized targeting. Also, the difference in the number of animals that responded to NAc core and shell stimulation likely underpins the difference in the population-based analysis described above -- the percent reduction in binge size was roughly equivalent in across stimulation sites in the individuals that responded.

### Experiment 3 - Behavioral and electrical characterization (without stimulation)

#### Correlating stimulation outcomes with reward-related behavior

It was our hypothesis that innate variation in NAc core and shell networks would be a common source of variation in reward-related behavior and stimulation outcomes. Thus, we expected to see a relationship between variation in reward-related tasks and stimulation outcomes. The behavioral metrics of the 12 included rats were grouped based on the rat’s individual response to stimulation as defined previously (R - responder and NR - non-responder for each stimulation target). Differences between R and NR groups were evaluated with t-tests and none of the behavioral measures showed a potential to differentiate the R group from the NR group for core- (Figure 4A) or shell- (Figure 4B) targeted stimulation.

**Figure 4.**
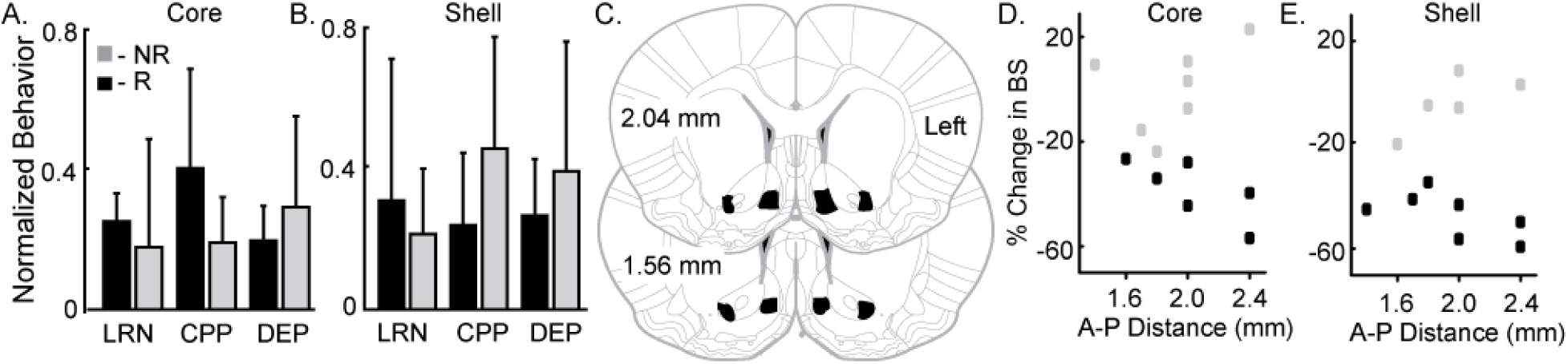
Variation in reward-related behavior and electrode location does not relate to stimulation outcomes. Normalized behavioral data grouped by core (**A**) and shell (**B**) DBS response type --responders (R; black) and non-responders (NR; grey). No significant differences were observed between R and NR groups for the following outcomes: 1) total distance travelled during locomotor response to novelty (LRN); 2) change in the percent of time spent in the initially non-perferred chamber during conditioned place preference (CPP); and 3) percentage increase in food intake after 24 hours of food deprivation (DEP). **C**. All 12 rats included in the analysis had electrode locations within the bilateral NAc core and shell with electrodes localized within the black shapes collapsed onto two representative coronal sections. The largest variation in electrode positioning occurred along the anterior-posterior (A-P) dimension (1.4 to 2.4 mm anterior to bregma) No qualitative relationship between electrode placement along the A-P axis in NAc core (**D**) or shell (**E**) corresponded to stimulation outcomes -- responder (black) or non-responder (grey).

#### Correlating stimulation outcomes with electrode localization

Figure 4C-E illustrates the relationship of anterior-posterior (A-P) position in the core (Figure 4D) and the shell (Figure 4E) and the corresponding stimulation outcomes (black --responders; grey -- non-responders). Variation of electrode location within the A-P dimension displayed no qualitative relationship with stimulation outcomes.

#### Correlating stimulation outcomes with local field potential activity

The lasso was able to use information contained within LFP features to determine which response group an animal belonged to with an average accuracy for core stimulation of 72% (standard deviation ± 5%), outperforming the models produced from random permutations of the data (49% accuracy ± 11%) with an effect size of 1.13 (Figure 5C). The lasso models classifying shell stimulation outcomes performed with an average accuracy of 65% (standard deviation ± 7%), outperforming the models produced from random permutations of the data (49% accuracy ± 11%) with an effect size of 1.05 (Figure 5E). Finally, each rat with a significant reduction in binge size was grouped by the target (NAc core or shell) that produced the largest average reduction in binge size across the three stimulation sessions. LFP features were able to match individual rats to the most effective target for stimulation using lasso with an average accuracy of 76% (standard deviation ± 7%) compared to 59% (standard deviation ± 8%) for the permuted data with an effect size of 0.96 (Figure 5D).

**Figure 5.**
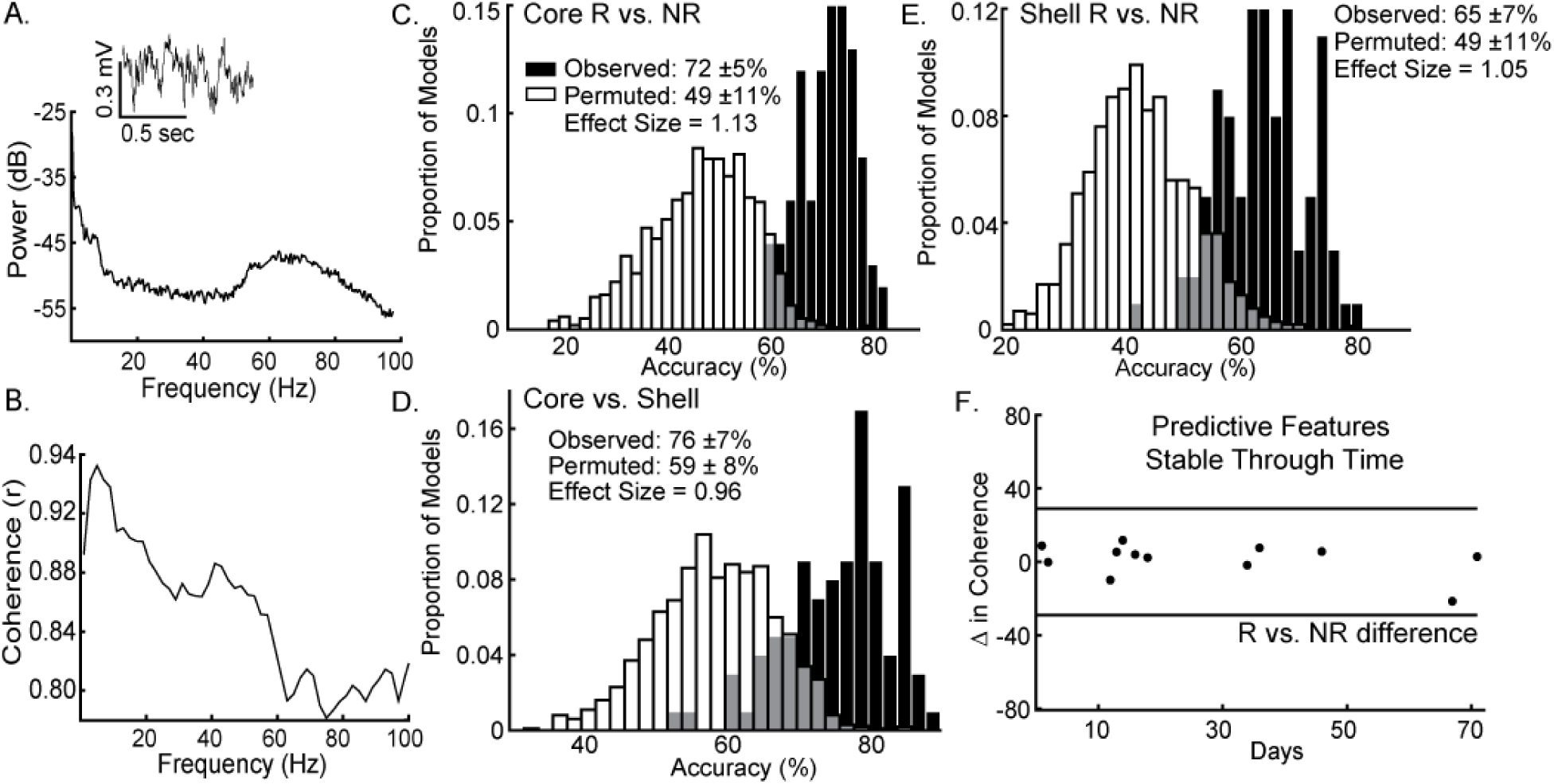
Local field potential (LFP) features recorded from ventral striatum can classify individual stimulation outcomes and are stable through time. **A**. Inset of a raw LFP trace from the left NAc core with its corresponding power spectral density plot. **B**. Corresponding coherence plot showing phase relationships across frequencies between the left NAc shell and right NAc core. The distribution of accuracies from classifying NAc core (**C**) and shell (**E**) stimulation responders (R) from non-responders (NR) using the observed data (black) and the permuted data (white) with mean accuracy ± standard deviation listed for each distribution. Effect sizes between observed and permuted distributions are also shown. **D**. Distribution of accuracies classifying the optimal target for stimulation (core vs. shell) for each animal using the observed data (black) or the permuted data (white). **F**. The difference in delta coherence (between the left NAc core and right NAc shell) from recording day T1 to T2 (up to 71 days apart) was smaller then the difference observed between the groups of animals that preferentially responded to core or shell.

It is important to note that each rat had 2 LFP recording sessions separated by up to 70 days and each recording session was separately incorporated into the model. Therefore, only LFP features that had stable differences between groups (e.g., R vs. NR) across time were selected and used by lasso. An example of one of the selected LFP features is shown in Figure 5F showing that the feature was less different between day 1 and day 71 within animals than observed between the response and non-response groups (Figure 5F -- black horizontal lines). This finding indicates that the information about stimulation outcomes extracted from LFP signals has stability through time.

To determine which components of the LFP signal contained information about stimulation outcomes, each feature’s univariate performance in logistic models (% accuracy) was compared to how commonly those features were included in the multivariate (lasso) models (% survival). Table 1 lists the top 5 LFP features from the logistic and lasso models of core and shell stimulation outcomes (R vs. NR) and the classification of the optimal target for each animal (core vs. shell). This exploration revealed a predominance of delta band features in the logistic models that did not translate to survival in the lasso models suggesting that while delta features contained the most information about outcomes, this information was likely highly redundant. Thus, only one delta feature tended to be included in the lasso models. Arrows in the table indicate the directionality of the feature differences between groups.

## Discussion

These experiments demonstrate that deep brain stimulation of either the nucleus accumbens core or shell, regions with known differences in brain connectivity and distinct functional roles in appetitive behaviors, have a similar capacity to alter “binge-like” feeding behavior. Experiment 1 demonstrated that despite titration across multiple stimulation parameters only subsets of animals show significant changes in binge behavior with stimulation in either of the tested targets. Experiment 2 confirmed this finding and an evaluation of individual responses across the first two experiments illustrates that 66% of rats only respond to DBS in one of the two targets, supporting the likelihood that personalized target selection could improve treatment outcomes. Experiment 3 demonstrated that variation in stimulation outcomes could be, in part, explained by individual differences in recorded local field potential activity using a machine learning-based approach (lasso). Most importantly, ventral striatal oscillations were also capable of classifying the most effective stimulation target for each individual, demonstrating the feasibility of using network activity to personalize target selection for neuromodulation-based treatments.

The translational relevance of this work is supported by previously observed treatment outcome variability in clinical studies of focal stimulation in disorders of appetitive behavior [5, 13, 40]. As an example, in a study using repetitive transcranial magnetic stimulation of the medial prefrontal cortex for patients with binge eating, differences in cortical-striatal network activity were shown to correlate with response to stimulation [27]. Therefore, it is notable in this study that a large proportion of animals that failed to respond to stimulation in one brain target (NAc shell), responded to stimulation in an alternative target (NAc core). Further, results from this study suggest that network activity recorded in the ventral striatum contains information that can predict the optimal target for stimulation on an individual basis. This finding suggests that even in this outbred rat model of binge eating there are likely individual differences in the networks contributing to the behavioral expression of binge eating.

The assertion that variation exists across individuals in the specific cortical-striatal networks that underpin the expression of appetitive behavior is supported by a rich literature including the well characterized spectrum of goal-directed to habitual behavior [41–44]. Thus, the striatal sub-regions driving binge like behavior could vary across individuals and impact which striatal target (NAc shell vs. core) is most likely to modulate binge behavior. Patients with binge eating have also been shown to display altered function in distinct networks including the reward/salience network [45–47] and/or the cortical control network [48–51] using non-invasive methods to assess network activity. Altered function of one of these networks may be enough to perpetuate binge eating [52], and our work in rats suggests that even within the ventral striatum, different sub-circuits (involving the NAc core or shell) may be differentially contributing to the maintenance of binge eating across rats. Taken together, clinical and pre-clinical studies suggest that a single stimulation target may not have the capacity to reduce binge eating across all individuals, and measures of relevant network activity could guide the selection of an effective stimulation target for each individual.

In order to translate personalized targeting of focal stimulation to patients, a non-invasive method of measuring network activity would be required prior to the intervention. Therefore, it is important to consider the relationship between information extracted from LFP oscillations recorded from depth electrodes reported in this study and non-invasive methods of measuring related network activity in patients. Our data suggest that inter-hemispheric coherence at low frequencies (delta and theta) may be a rich source of information about DBS outcomes. Previous work has established that correlation exists between these LFP features and fMRI derived measures, including resting state functional connectivity [53–55]. The work presented in this study supports the inclusion of the ventral striatum and interconnected cortical regions for future investigations that attempt to use brain activity to guide targeting of focal stimulation for binge eating and related disorders of appetitive behavior.

Overall this study was limited by the scope of information used (recordings from bilateral NAc core and shell) to build our predictive models. Thus, increasing the number of recording sites to include regions in the distributed feeding circuit (e.g., hypothalamic/brainstem, medial prefrontal and orbitofrontal cortex) would be important for future studies. In particular, recording from cortical regions would have translational relevance to non-invasive clinical measures of brain activity (e.g., EEG) in addition to MRI derived features. Future studies will incorporate pre-stimulation recordings in order to capture network dynamics in treatment naïve animals. In addition, although using a penalized regression (lasso) mitigated the problem of having many more predictor variables than observations, a larger sample size would allow for testing of the tuned multivariate regressions on naïve datasets and provide more power to correlate variation in electrode location with stimulation outcomes. We cannot rule out the possibility that variation in targeting within the NAc sub-regions also contributed to stimulation outcome variation. Inclusion of a female cohort would have increased the generalizability of this study as more women suffer from binge eating compared to men. Lastly, while none of the reward-related behaviors tested in this study showed the potential to predict stimulation outcomes, additional behaviors with differential NAc core and shell dependence may better characterize the individual variation that may underlie the variation in stimulation outcomes [43, 56].

## Conclusion

For the treatment of many psychiatric disorders, as demonstrated here in a rat model of binge eating, a single target for focal stimulation may not be effective across all individuals. Rather, an individualized treatment approach that uses network activity to guide personalized target selection could lead to improved outcomes for neuromodulation-based treatments.

